# Diffusion-based artificial genomes and their usefulness for local ancestry inference

**DOI:** 10.1101/2024.10.28.620648

**Authors:** Antoine Szatkownik, Léo Planche, Maïwen Demeulle, Titouan Chambe, María C. Ávila-Arcos, Emilia Huerta-Sanchez, Cyril Furtlehner, Guillaume Charpiat, Flora Jay, Burak Yelmen

## Abstract

The creation of synthetic data through generative modeling has emerged as a significant area of research in genomics, offering versatile applications from tailoring functional sequences with specific attributes to generating high-quality, privacy-preserving in silico genomes. Notwithstanding these advancements, a key challenge remains: while some methods exist to evaluate artificially generated genomic data, comprehensive tools to assess its usefulness are still limited. To tackle this issue and present a promising use case, we test artificial genomes within the framework of population genetics and local ancestry inference (LAI).

Building on previous work in deep generative modeling for genomics, we introduce a novel, frugal diffusion model and show that it produces high-quality genomic data. We then assess the performance of a downstream machine learning LAI model trained on composite datasets comprising both real and/or synthetic data. Our findings reveal that the LAI model achieves comparable performance when trained exclusively on real data versus high-quality synthetic data. Moreover, we highlight how data augmentation using high-quality artificial genomes significantly benefits the LAI model, particularly when real data is limited. Finally, we compare the conventional use of a single synthetic dataset to a robust ensemble approach, wherein multiple LAI models are trained on diverse synthetic datasets, and their predictions are aggregated.

Our study highlights the potential of frugal diffusion-based generative models and synthetic data integration in genomics. This approach could improve fair representation across populations by overcoming data accessibility challenges, while ensuring the reliability of genomic analyses conducted on artificial data.

## 1 Introduction

Generative AI has been widely adopted across various domains thanks to recent advances in novel algorithms and architectures. Synthetic outputs generated by these models often capture essential characteristics of real-world data and, in some cases, can match human production in specific tasks such as question-answering (Ke et al., 2024). Synthetic data is also useful across fields, as it enables the creation of diverse datasets that can improve model training, facilitate testing in resource-limited areas, and help overcome data scarcity (Beaulieu-Jones et al., 2019; Lu et al., 2023). In genomics, generated data is particularly valuable because it could allow for the study of genetic diversity without compromising individual privacy and enable the testing and application of genome-wide-association-studies or other methodological developments without the need for exhaustive real-world samples (Yelmen and Jay, 2023; Wharrie et al., 2023).

However, compared to image and text domains, artificial genomic data is substantially more challenging to assess both in terms of quality and utility, due to the complex and very high dimensional data structure shaped by billions of years of evolution. Here, we define the utility of artificial genomes for a given task based on whether a model solving the downstream task achieves comparable performance using synthetic data as with real data. In this context, predictive performance in various tasks can serve as an indicator of utility while population genetics summary statistics as an indicator of quality (Yelmen et al., 2021, 2023; Szatkownik et al., 2024a,b), albeit these two concepts overlap in most cases (Lacan et al., 2023, 2024; Montserrat et al., 2019). Previous studies on the utility of synthetic genomic data have explored various aspects, including genomic imputation, detection of natural selection, and inference of evolutionary parameters (Yelmen et al., 2021; Wang et al., 2021). In this work, we focus on one such aspect, local ancestry inference (LAI), extending the work of Montserrat et al. (2019).

The genetic material of individuals can be inherited by ancestors coming from different populations. Each individual in a population might have different ancestry coefficients, representing the proportions of their genome derived from multiple ancestral gene pools due to admixture events (Frichot et al., 2014). Upon closer examination, one can observe that these ancestries are distributed across the genome. The objective of LAI methods is to determine which part of the individual’s genome comes from which ancestral population, allowing refined analysis of past demographic events and natural selection with respect to these specific ancestries. Therefore, ancestral reference genomes used in LAI should not only be representative of correct genome-wide global structure with longrange dependencies, but also have to conditionally represent the correct local structure such as linkage between genomic positions. Many algorithms for LAI have been developed over the years, traditionally using Hidden Markov Models (Price et al., 2009; Salter-Townshend and Myers, 2019; Lawson et al., 2012; Skov et al., 2018; Planche et al., 2024), statistical methods (Yang et al., 2013), PCA (Brisbin et al., 2012), and more recently, graph optimization (Dias-Alves et al., 2018; Wei et al., 2023) and machine learning (Maples et al., 2013; Montserrat et al., 2020; Hilmarsson et al., 2021).

In this study, we introduce a novel frugal latent diffusion model for human genetic data. Previous works in artificial genome generation have utilized restricted Boltzmann machines (RBMs), variational autoencoders (VAEs), generative adversarial networks (GANs), generative moment-matching networks (GMMNs), probabilistic circuits (PCs) and large language models (LLMs) (Yelmen et al., 2021; Perera et al., 2022; Dang et al., 2023; Zhang et al., 2024), see Yelmen and Jay (2023) for a review. Some of these models were designed to be conditional on population labels, other demographic parameters, or the preceding genomic region, which permits refining the learned data distribution and conditional sampling of generated genomes based on these labels (Montserrat et al., 2019; Booker et al., 2023; Yelmen et al., 2023). Denoising diffusion probabilistic models (DDPM) have been originally tested for image and audio generation and are known to be useful for tasks involving high-dimensional data synthesis (Sohl-Dickstein et al., 2015; Ho et al., 2020). Their applications range from image super-resolution, text-to-image generation, inpainting of images or computational fluid dynamics data to molecular structure generation (Kong et al., 2021; Dhariwal and Nichol, 2021; Ramesh et al., 2021; Rombach et al., 2022; Saharia et al., 2023; Lugmayr et al., 2022; Shu et al., 2023; Hoogeboom et al., 2022; Xu et al., 2022). Recently they have been applied in functional genomics for generating short DNA sequences conditioned on species type, expression level, or cell type with success (Li et al., 2023, 2024; Pinello, 2024). Yet, to our knowledge, no diffusion model has been designed for population genetics, with a focus on capturing genomic variation and population genetic diversity. In this work, we aim to leverage the relative ease of training diffusion models (especially compared to GANs) to develop a DDPM conditioned on continental group labels (also called super-populations).

We trained our frugal diffusion model, Light PCA-DDPM, on a diverse panel of human genomic data and examined the synthetic data generated. After evaluating the quality of these artificial genomes in terms of summary statistics, we assessed their usefulness for LAI. For that, we relied on a recent deep learning based method called LAI-Net (Montserrat et al., 2020), due to its high performance and low time complexity. We trained LAI-Net on this synthetic data and compared predictive performance to LAI-Net trained on real data. We further increased synthetic sample size and studied the impact on LAI performance. We also explored a diffusion-based data augmentation scenario by adding synthetic data to training sets including varying amount of real samples, mimicking different data accessibility scenarios. Finally, we investigated a Deep Generative Ensemble approach (DGE) (van Breugel et al., 2023), that involved the training of multiple generative models and LAI-Nets to mitigate potential generative errors in downstream tasks.

## 2 Materials and Methods

### 2.1 Dataset description

#### Full dataset

The 1000 Genomes (1KG) Project is a database of human genome sequences, designed to capture the broad genetic diversity of the human species by sampling individuals from 26 distinct populations scattered across the continents (The 1000 Genomes Project Consortium, 2010). Each population is represented by a number of haplotypes ranging from 122 to 226 (**FIG.S1**). The dataset used in this study is derived from the one curated by Yelmen et al. (2023), that included 2,504 individuals, corresponding to 5,008 phased haplotypes (i.e., one haplotype inherited from each parent per individual). It comprises 65,535 contiguous SNPs spanning chr1:534247-81813279 (approximately 80 megabase pairs) within the Omni 2.5 genotyping array framework. The data is organized as follows: rows represent phased haplotypes, while columns indicate allele positions, coded as 0 if the nucleotide matches the reference genome (GRCh37), and 1 if it represents a point mutation. This results in a binary matrix representation of the dataset.

#### Three-continent dataset

A subset of the Full dataset targeting unadmixed individuals classically used to create LAI reference panels, i.e. individuals labeled as African (AFR) (removing ASW and ACB as these are admixed), East Asian (EAS) and European (EUR), amounting to 1,510 samples. We applied a train-test split of 80-20%, resulting in 1,208 and 302 diploid sequences respectively. The training set was further sub-sampled to an equal amount of 396 diploid sequences per ancestry (*i*.*e*., 1,188 individuals in total).

#### Haptool-admixed test sets

The remaining 302 sequences were used as input data for the admixture forward-simulator Haptools (Massarat et al., 2023) to produce a final amount of two sets of 156 admixed individuals for test data (*i*.*e*., 312 haplotypes in total), see section 2.3 for details.

#### AMR test set

A subset of the Full dataset including individuals labeled as American (AMR) (CLM, MXL, PEL, PUR, ACB, ASW), amounting to 504 diploid sequences.

### 2.2 LAI-Net

LAI-Net (Montserrat et al., 2020) is a supervised algorithm that takes as input phased diploid sequences. These haplotypes are then split into non-overlapping SNP windows for which the size is a user-defined parameter and represents the resolution of the algorithm. For each of these windows a neural network is assigned, which outputs a vector of probabilities across the ancestral populations. A second layer, consisting of learned convolutions, refines the inference by smoothing the prediction of a window given neighboring windows. In our application, we fixed the window size to 500 SNPs and maximum number of training epochs to 700. The ancestral sources used were Africa (AFR), Europe (EUR), and East Asia (EAS). All other parameters were default.

### 2.3 Local ancestry inference

We trained multiple LAI-Nets on both real (Three-continent dataset) and synthetic datasets, each comprising an equal number of EUR, AFR, and EAS ancestral samples. The synthetic data were generated using a conditional diffusion model (see Subsection.2.4) and a baseline random matrix model (see Subsection. 2.5).

For model evaluation, we used both real haplotypes (AMR test set) and artificially admixed haplotypes with ground-truth admixture information as target sequences (haptool-admixed test sets). The AMR test set consisted of 504 admixed American samples. We used separate held-out samples (i.e., not used as ancestral samples for training LAI-Net) from the three ancestral sources to construct the haptool-admixed test set using the forward simulator Haptools (Massarat et al., 2023). The simulator takes as input 1KG project haplotypes from the ancestral populations and a set of parameters: the ancestry coefficients (i.e., percentage of ancestries over the whole genome) desired for the output sequences and the number of generations under which the simulation takes place. Then the sequences are shuffled based on these parameters and recombination is simulated to produce admixed haplotypes. This process allows us to have full control over the admixture event and know the ancestry at each genomic site. We ran the simulator to admix EUR, AFR and EAS held-out samples with ancestry coefficients of 0.51, 0.16 and 0.33 respectively, which could correspond to estimates for some real American populations when EAS is considered as a proxy for indigenous AMR component (Medina-Muñoz et al., 2023; Browning et al., 2018). The number of generations since admixture was fixed at 30 (haptool-admixed-30) and 60 (haptool-admixed-60). There is a relation between the number of generations and the size of genomic windows defined by crossover events (See Appendix.S1.3). The number of generations was chosen to have an easy task and a hard one, both with the property that window sizes are larger than the resolution of LAI-Net (set to 500 SNPs), thus avoiding impractical data.

Overall, we checked how accurately each ancestry was assigned by computing per ancestry, whether the prediction for a given window was correct or not. For simulated test data with ground truth, the label of a window was set as the dominant ancestry in that window. The accuracy per ancestry was averaged over all windows and all samples. We either kept the results per ancestry or did a weighted average where the weights correspond to the true ancestry coefficients. For AMR test data, since true ancestral compositions are not available, we assessed the concordance across the genome (identity percentage) between LAI-Net trained with real and synthetic ancestral samples.

### 2.4 Light PCA-DDPM

A Denoising Diffusion Probabilistic Model (DDPM) (Sohl-Dickstein et al., 2015; Ho et al., 2020) consists of a denoising neural network trained to predict the noise of a corrupted sample or the original sample. In practice it was shown to work best by predicting the noise (Ho et al., 2020). The corruption phase, called forward process, takes a sample *x*_0_ ∈ ℝ^*D*^ from the real data distribution, where *D* is the dimension of the data space, and creates a sequence of increasingly noisy versions: *x*_1_, *x*_2_, … *x*_*T*_ ∈ ℝ^*D*^. This sequence is produced by gradually adding Gaussian noise to the data according to some noise schedule *β*_1_, … , *β*_*T*_ , *i*.*e*.,

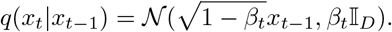

This forward noising operation can be computed in closed-form, allowing one to bypass the sequential process of gradual noising and landing directly at the desired time *t, i*.*e*.,

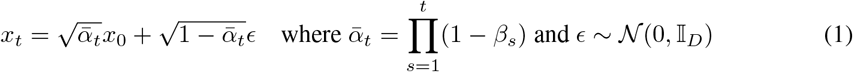

The noise level *β*_*t*_ increases linearly with time, from 0.0001 to 0.02. The final *x*_*T*_ is an isotropic Gaussian noise sample.

While the forward process does not involve any optimization, the backward process entails inverting the forward noising procedure by training a neural network *f*_*θ*_ to progressively denoise each sample *x*_*t*_ in decreasing order of *t, i*.*e*., from *t* = *T* to *t* = 0. In practice, to train a diffusion model, we:

1. Sample *x*_0_ from the dataset, sample a random time step *t* and sample random Gaussian noise *ϵ*
2. Corrupt the data sample *x*_0_ into noisy sample *x*_*t*_ via the forward process Eq. (1)
3. Predict the noise in *x*_*t*_ by running it through *f*_*θ*_
4. Minimize the loss 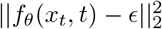 with respect to *θ*, where the loss is the MSE between the true noise and predicted noise, *i*.*e*. the goal of the network is to predict *ϵ* from a noisy observation, conditionally on time step *t*.

Once the model is trained, the backward operation corresponds to generative modeling. To generate new fake data, we sample from an isotropic Gaussian distribution, and iteratively remove the noise incrementally by going over all the time steps, from *T* to 0:

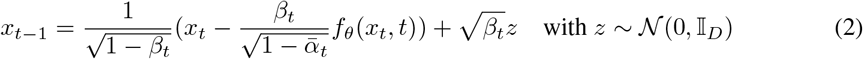

For our implementation, Light PCA-DDPM, we adopted a similar approach to that of Light PCA-WGAN (Szatkownik et al., 2024b) to maintain low complexity. Here, instead of a WGAN, we employed a DDPM and additionally conditioned the denoising neural network on the super-population from which an original sample *x*_0_ comes from, *i*.*e*., AFR, EUR or EAS. Light PCA-DDPM combines an initial PCA on real data, retaining all principal components (PCs), with generative modeling in that latent space (**FIG. 1**). The resulting PC scores are split into a high variance (first six dimensions/ PCs) and low variance (remaining 2134 dimensions / PCs) parts. The DDPM is trained on the high variance PCs which display multi-modal structure. Then a multivariate normal distribution (MVN) is constructed to model the remaining 2134 PCs. This MVN is parameterized by a mean vector 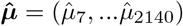 extracted by taking the mean over the low variance real PC scores across all individuals, and a diagonal covariance matrix 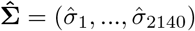 computed similarly using the low variance real PC scores across all individuals.

**Figure 1.**
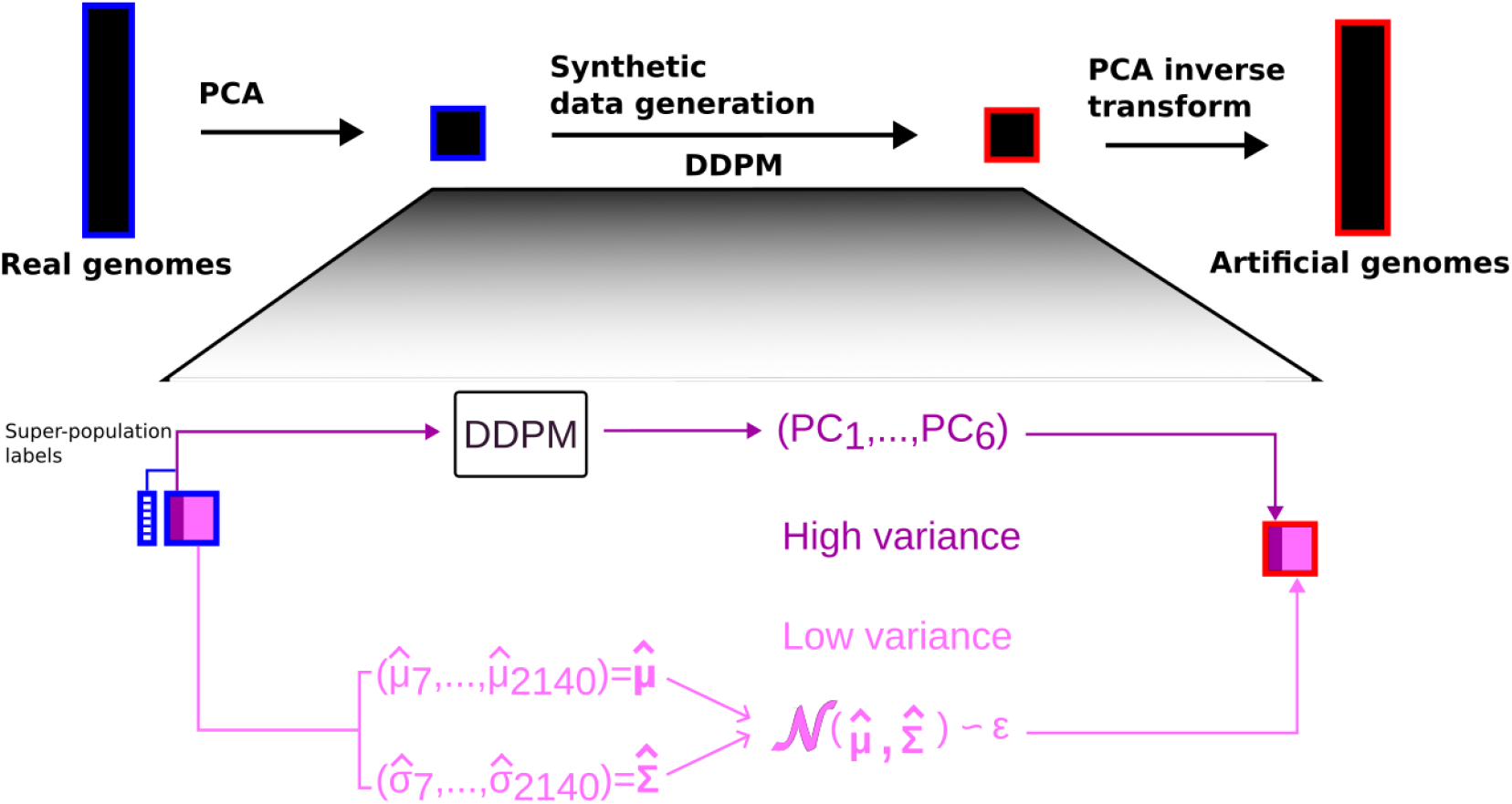
PCA-DDPM. PCA is first applied to real SNP data. During training, only the DDPM is involved, learning to mimic the distribution of real data in the latent space, conditioned on the ancestral super-population labels (matrices in PCA space are square matrices). At sampling, the high variance PC scores (dark pink) are generated by the DDPM, while the low variance PC scores (light pink) are sampled independently from the MVN. The two components are then concatenated, inverse transformed using inverse PCA, and binarized to produce artificial genomes.

For generation, we sample random Gaussian noise and a super-population label, which are processed through the denoising network to produce the high-variance PC scores for the desired ancestry. The corresponding low variance PC scores are generated by sampling the MVN independently of the high variance PC scores and unconditionally to the super-population label. Finally, the high and low variance synthetic PC scores are concatenated and inverse transformed back into SNP space with PCA inverse transform. The resulting sequence is then binarized with fixed threshold set to 0.5.

Our implementation of *f*_*θ*_’s architecture is detailed in Appendix.S1.2. We will hereafter use the term *PCA-DDPM* when referring to *Light PCA-DDPM*, for the sake of simplicity.

### 2.5 Bernoulli model

We also designed a simple generative model for SNP data based on one-point statistics only, to compare the artificial genomes against. This baseline random matrix model is a mixture of three Bernoulli distributions, with parameters derived from the allelic frequencies at each site in the AFR, EUR, and EAS super-populations of the real data, respectively. More precisely, for each site and group, we computed the frequency *f* of the alternative allele in the reference individuals. For each group, we then generated artificial genomes by randomly selecting the alternative allele at each site with probability *f*. We will hereafter use the term *Bernoulli model*, when referring to this mixture of three Bernoulli distributions.

### 2.6 Deep Generative Ensemble

In real-world applications of generative AI, the most straightforward approach that balances both privacy and usability is for the data holder to train a single generative model to produce synthetic data, which is then shared (*Naive* scenario). The downstream data user can then utilize this synthetic dataset, for instance, to train a machine learning (ML) model. However, previous work shows that treating synthetic data as equivalent to real data leads to poor downstream model training and evaluation, with higher susceptibility to perform worst for under-represented classes (van Breugel et al., 2023). Deep generative ensemble (DGE) was proposed to avoid these problems. In this framework, the data holder trains *K* generative models (*K >* 1, and using different seeds) and generates *K* synthetic datasets which are shared. The data user trains *K* ML models with these synthetic datasets and aggregates the results by averaging the predictions. Additionally to these previous scenarios presented in (van Breugel et al., 2023), we explored two novel ones, the *Naive big* and *DGE big* scenarios. The *Naive big* refers to having larger synthetic sample size than *Naive* (here we set it to be four times larger than real sample size), while the *DGE big* scenario, refers to the *DGE* setup with synthetic data being four times larger than real sample size. Here we fixed *K* = 5 and used PCA-DDPM as the generative model. Moreover, each scenario was repeated 10 times to have an estimation of the variance.

## 3 Results

### 3.1 Quality of artificial genomes

We first assessed the capacity of PCA-DDPM (Methods. 2.4) to generate highly diverse and high quality artificial genomes. The quality of the synthetic data was evaluated via a set of population genetics summary statistics and compared to the previously established Light PCA-WGAN model (Szatkownik et al., 2024b) and to the baseline Bernoulli model. All models were trained on the Full dataset of 2,504 individuals spread across the five super-populations. PCA-DDPM performed on par with Light PCA-WGAN on summary statistics measuring global patterns of genetic variation. Notably, they retrieved population genetic structure as captured by the first axes of a PCA and the 1D Wasserstein distance between PC scores was slightly better for PCA-DDPM than for PCA-WGAN. Interestingly, the Bernoulli model outperformed both of these models on the first two PCs (**FIG. S2**). This result can be attributed to the modes of the PC scores for the Bernoulli model being closely aligned with those of the real data, albeit having a substantially shrunk distribution (**FIG.S3**), demonstrating that the Wasserstein distance can be very sensitive to mode densities as reported in the literature (Montavon et al., 2016). Furthermore, PCA-DDPM and PCA-WGAN also had good and similar abilities to capture allelic frequencies. Since the Bernoulli model was explicitly given allelic frequencies, it also performed well on this metric as expected. Both PCA-DDPM and PCA-WGAN exhibited comparable local haplotypic structures based on linkage disequilibrium (LD) measurements (see **FIG. S2**), while the Bernoulli model, designed to capture only one-point statistics, was unable to capture the LD structure. Finally, PCA-DDPM and PCA-WGAN again displayed similar behavior on the three-point correlation statistics (**FIG. S2**). However, while the Bernoulli model showed low correlation in short-distance three-point interactions, it performed relatively well on the long-distance ones. This could be explained by the fact that short-range interactions among SNPs are primarily influenced by physical linkage, while long-range interactions are largely determined by population structure. Consequently, the Bernoulli model, which is fitting each super-population independently, more effectively captures long-range correlations arising from differing allele frequencies across super-populations (see also inset in LD plot of **FIG.S2**).

### 3.2 Ancestry inference on ground truth data

LAI-Net trained on real AFR, EAS, EUR individuals (Three-continent dataset) and tested on the haptool-admixed test sets yielded good overall accuracy (see Table **1**). Synthetic data generated by PCA-DDPM trained on the Three-continent dataset performed similarly to real data (though with slightly lower overall accuracy), suggesting that the synthetic dataset may contain information content comparable to that of the real dataset. Furthermore, even though LAI-Net trained on synthetic data coming from the Bernoulli model performed surprisingly well, the accuracy was below that of the other classifiers by ∼ 10%. This suggests, as expected, that allele frequency information alone is not sufficient for LAI and that our diffusion model retain characteristics that are relevant for this task.

### 3.3 Ancestry inference on admixed American samples

We further wanted to test the utility of artificial genomes when performing LAI on real admixed populations. Since true ancestral composition of real genomes are not available, we instead computed the percentage of identity between predictions on admixed American populations (AMR test set) from LAI-Net trained on real data and LAI-Net trained on synthetic data generated via PCA-DDPM or the Bernoulli model. For the pairs of predictions coming from real data & synthetic data from PCA-DDPM, we achieved 93% of identity over all populations, while with real data & synthetic data from the Bernoulli model, we achieved 83% of identity. Similar to the findings from the forward simulations, this suggests that artificial genomes generated via PCA-DDPM have richer information content than those generated via the baseline Bernoulli model.

We further investigated the identity percentages at the population level (**FIG. 2**). For both PCA-DDPM and Bernoulli model, the identity percentages were lowest for Mexicans (MXL) and Peruvians (PEL), which are two populations with the highest estimates of East Asian (EAS) ancestry. Additionally, LAI-Net trained on synthetic data predicted a higher proportion of EAS compared to when it was trained on real data. This tendency was more pronounced when LAI-Net was trained on synthetic data generated from the Bernoulli model.

**Figure 2.**
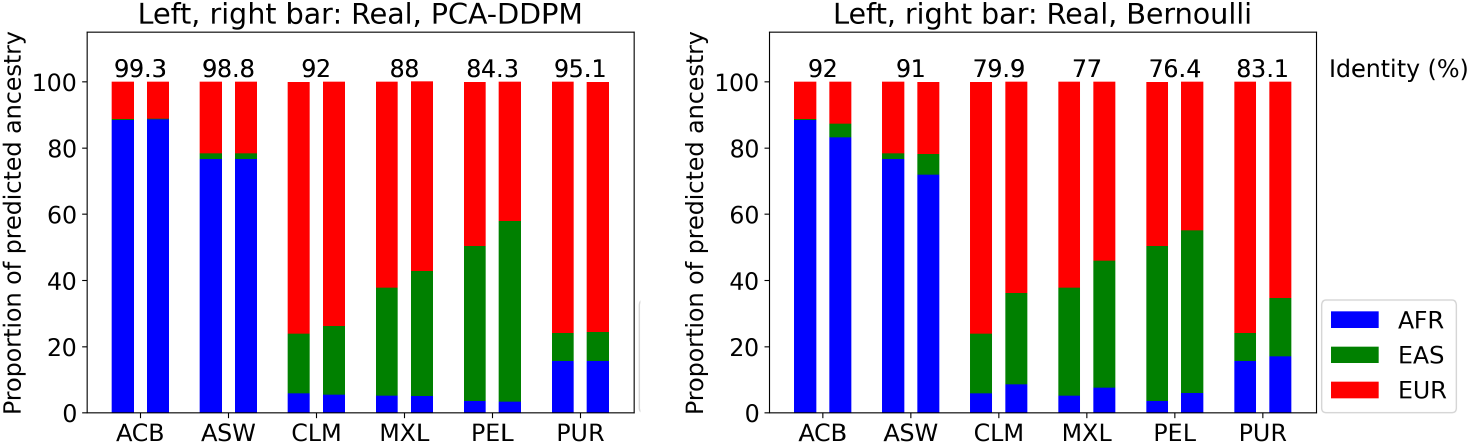
Identity (%) between predictions on AMR test set of LAI-Net trained on real and synthetic data. LEFT. Synthetic data is generated from PCA-DDPM. For each *x*, the left bar is the result of LAI-Net trained on real data and the right bar is the result of LAI-Net trained on synthetic data. The colors represent the proportion of predicted ancestry; AFR (blue), EAS (green), EUR (red); unfolded over populations within the American continent. The percentages of identity between predictions of both models are shown for each population on top of the bars. **RIGHT**. Synthetic data is generated from the Bernoulli model.

### 3.4 Increasing synthetic sample size

To test whether larger synthetic datasets can improve LAI, we generated synthetic data from both PCA-DDPM and Bernoulli model, with sample sizes being 2×, 4× or 6× larger than real data (**Table 2**). With increased sample sizes, we observed almost no performance improvement for LAI-Net trained on synthetic data from PCA-DDPM, except for a slight increase of 1.3% in accuracy for EAS predictions, hence we did not carry out the extensive computations required to test the remaining sample sizes. Similarly, training LAI-Net on synthetic data from the Bernoulli model with increasing sizes did not improve accuracy, but instead very slightly decreased it by ∼ 1% and lead to lower variances for 2*×* and 4*×* larger datasets.

### 3.5 Data augmentation

Synthetic data can be particularly useful to alleviate overfitting caused by small sample sizes. They can be used to augment the entire distribution of real data or specifically target low-density regions associated with under-represented classes. To investigate this, we trained LAI-Net on varying amount of real samples, ranging from 1% (= 12 haploid sequences) of the real training set to 100% (= 2136 haploid sequences), while having a fixed amount synthetic sequences (set to 8554). Each time, the generative model was trained on 100% of the real training data. This represents a scenario where a private data-holder shares either a generative model trained on private training data, or artificial genomes from one or multiple such generative models, but not the private data itself. For both haptool-admixed-30 and haptool-admixed-60 test datasets, LAI-Net trained on real data augmented with synthetic samples from PCA-DDPM outperformed other approaches when the percentage of true samples was low (up to 25 % of the original dataset). This indicates that data augmentation with high quality synthetic samples is beneficial for LAI when training data is limited and does not comprehensively represent the overall real distribution. Moreover, we also observed that the Bernoulli-augmented model performs worse than the non-augmented baseline for all percentages of true samples above 1%, which suggests that not all augmentation is good and adding data from a poor generative model can be detrimental to the classifier (**FIG.3**).

**Figure 3.**
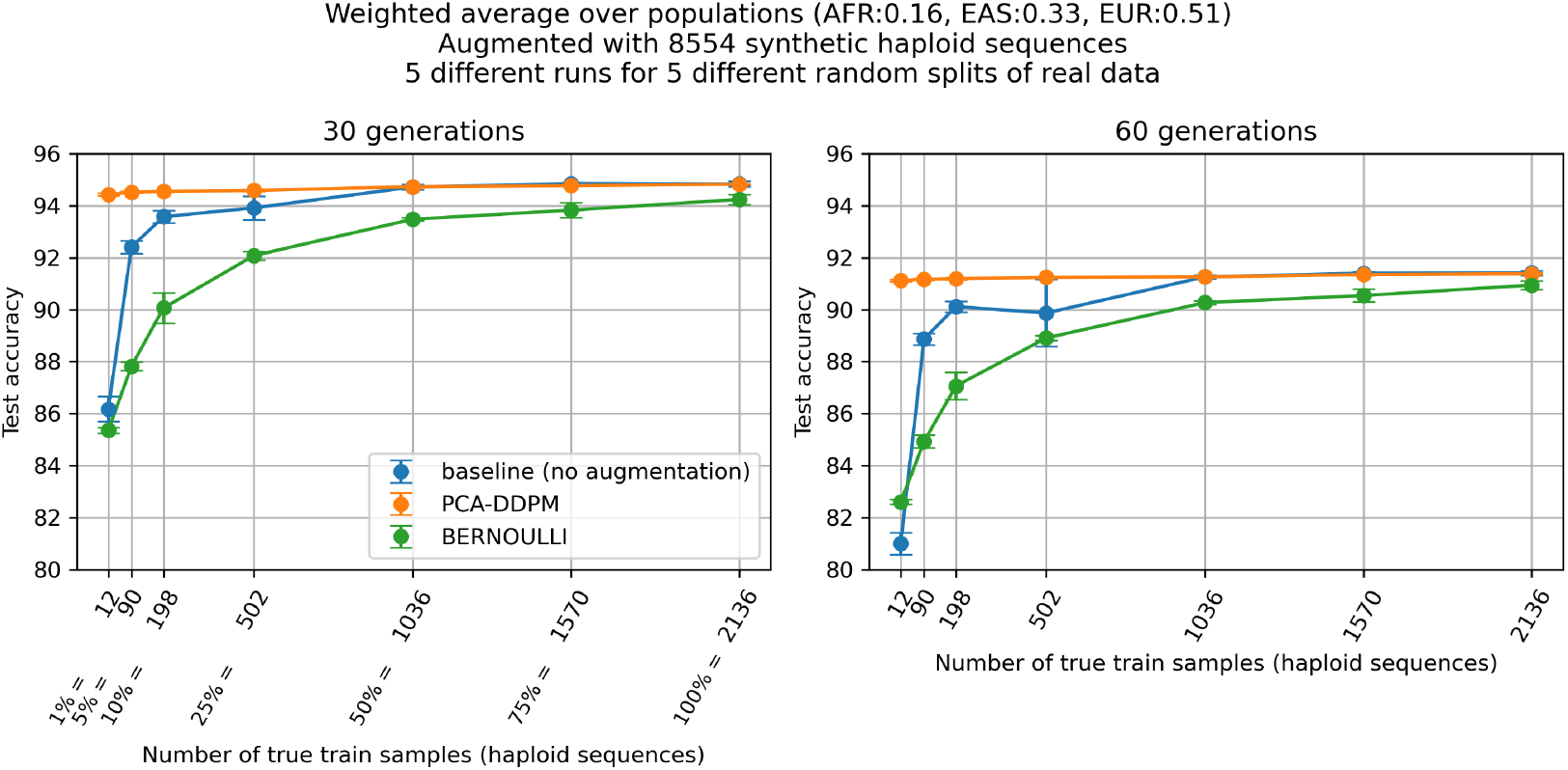
Data augmentation. Accuracy of LAI-Net evaluated on artificially haptool-admixed-30 (30 generations) and haptool-admixed-60 (60 generations) test datasets consisting of real AFR, EAS and EUR, and trained on: (i) varying amount of true training samples (blue curve); (ii) varying amount of true training samples augmented with 8554 synthetic haploid sequences from PCA-DDPM (orange curve).; (iii) varying amount of true training samples augmented with 8554 synthetic haploid sequences from the Bernoulli model (green curve). For each amount of true training samples, the experiment was repeated 5 times with different random splits of real data each time. The test accuracy was averaged over the populations with weights corresponding to the ancestry coefficients (AFR:0.16, EAS:0.33, EUR:0.51).

### 3.6 Deep Generative Ensemble

All different setups related to the DGE analysis (*Naive, Naive big, DGE* and *DGE big*, see Methods 2.6) performed only slightly worse than the original model trained on the real dataset (orange diamonds), with a mean accuracy of 94.84 ± 0.1, 94.16 ± 0.06, 94.42 ± 0.03, 94.23 ± 0.03 and 94.43 ± 0.03 for *Real, Naive, Naive big, DGE* and *DGE big*, respectively (**FIG. 4**). This corresponds to accuracies 0.4-0.7% lower for haptool-admixed-30 test data and 0.3-0.6% lower for haptool-admixed-60 test data. We observed that increasing synthetic sample size improved the model evaluation, i.e., *Naive big* yielded 0.2% improvement over *Naive*, and likewise *DGE big* yielded 0.2% over *DGE. DGE* was marginally above the *Naive* approach for haptool-admixed-30 test data, and comparable on haptool-admixed-60 test data, while *DGE big* and *Naive big* were comparable across both datasets. All these methods fall in between the test accuracy of a model trained on real data with sample size ranging from 25% to 50% of the real sample size (horizontal dashed lines in **FIG. 4**). Overall, considering y-scale of **FIG. 4**, the *DGE* approaches yielded marginal improvements over the *Naive* approach. Moreover, increasing synthetic sample size (*Naive big* and *DGE big*) only slightly increased test accuracy.

**Figure 4.**
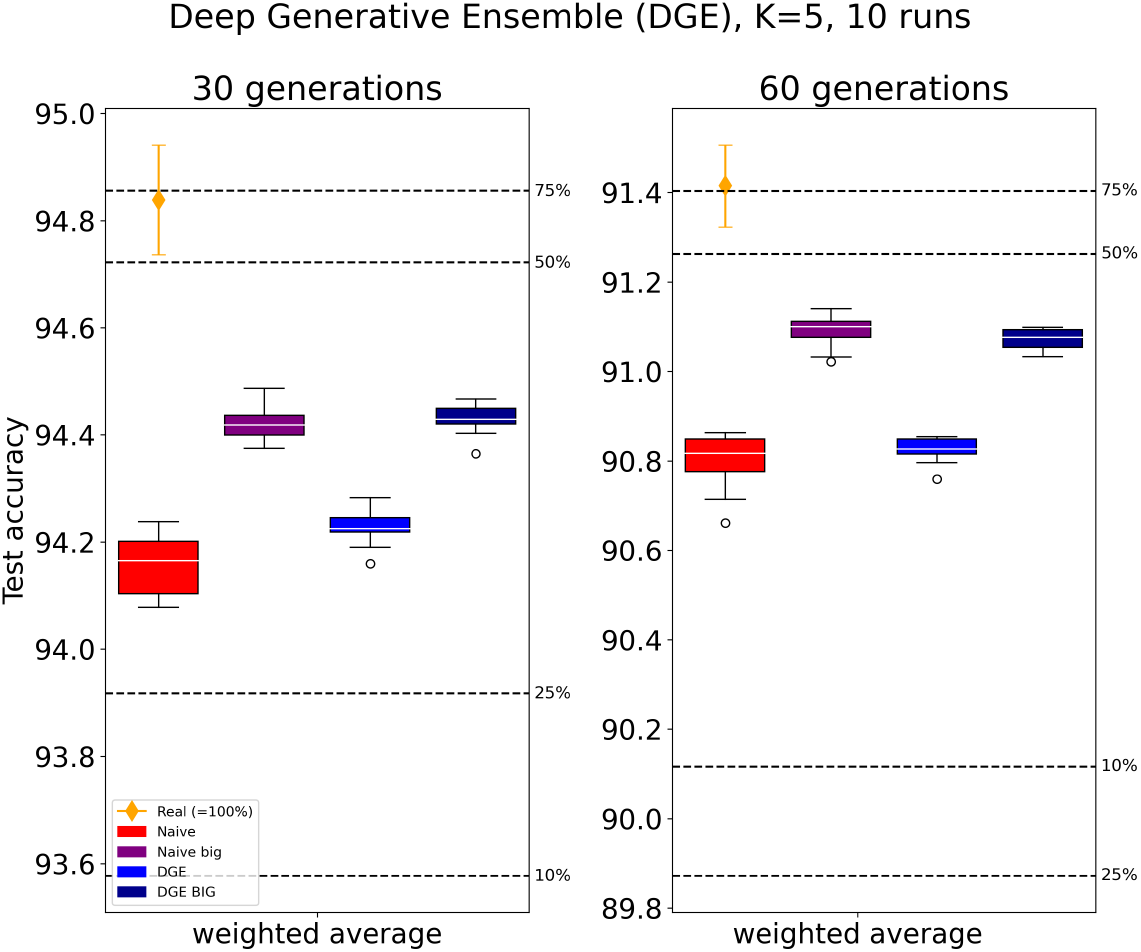
Deep Generative Ensemble. Accuracy of LAI-Net trained on only real data (orange diamond) or synthetic data from PCA-DDPM (Naive in red; Naive big in purple; DGE in blue; DGE big in dark blue). For each scenarios, we repeated the training of LAI-Net 10 times, and in particular for the *DGE* and *DGE big* (see Methods 2.6), the ensemble approach involved *K* = 5 iterations of training amounting to a total of 50 trained models. These results were averaged over *K*. The horizontal dashed lines correspond to LAI-Net trained on *x*% of real sample size (as in **FIG.3**.)

## 4 Discussion and conclusion

This study tackled the assessment of the usefulness of artificial genomes via a population genetics downstream task, namely local ancestry inference (LAI). We first developed a novel conditional diffusion model, Light PCA-DDPM, to generate high-quality synthetic haplotypes conditioned on super-population labels. Light PCA-DDPM, with only 400K learnable parameters, can be considered as an effective frugal alternative to more complex sequence modeling architectures while allowing the rapid generation of realistic synthetic data (see computational efficiency in Table S1 and quality in **FIG. S2**). We then benchmarked the generated genomes by training a neural network based LAI method, LAI-Net, on this synthetic data and evaluated its performance on datasets of varying degrees of difficulties. We observed similar test accuracy for LAI-Net trained on real data and on high-quality synthetic data, indicating that the information content, as perceived by a LAI method, is similar in synthetic and real data. In contrast, LAI-Net trained on synthetic data from the simple Bernoulli model displayed significantly worst performances with lower classification accuracies (Table 1). This demonstrates that higher-order information, as captured by deep generative models, are needed for LAI. Furtermore, even though increasing high-quality synthetic sample size did not affect LAI-Net performance, PCA-DDPM-based data augmentation was especially beneficial when real data was limited to the user, presenting a promising finding for practical applications where the complete distribution of real data is often not fully available through public databases. We also explored the Deep Generative Ensemble (DGE) approach which showed only marginal improvements over Naive scenarios. van Breugel et al. (2023) found that DGE does not always improve predictive accuracy over using real data, and the performance gain depends on the dataset and task to be solved. Training multiple classifiers in an ensemble fashion provides both the mean and variance of predictions, enabling the estimation of predictive uncertainty (Abdar et al., 2021). Beyond that, an important advantage of DGE is its ability to mitigate potential generative errors in downstream tasks by accounting for uncertainties in the generative model, and consequently, in the synthetic data (Decruyenaere et al., 2024; Räisä et al., 2023).

**Table 1:**
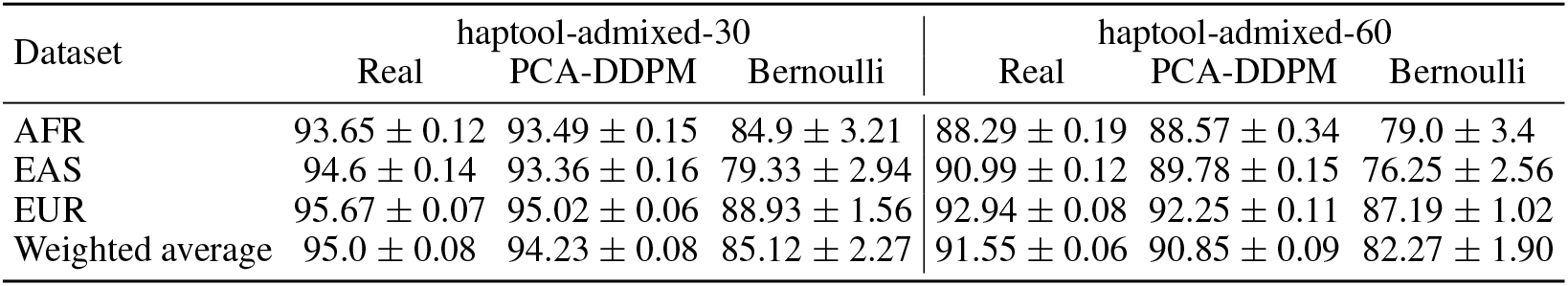
Accuracy (Precision in %) of LAI-Net trained on real or synthetic data and evaluated on haptool-admixed-30 and haptool-admixed-60 test datasets. Synthetic data is either coming from PCA-DDPM or Bernoulli model. Ancestry coefficients in test data are 0.16, 0.33, 0.51 for AFR, EAS, EUR respectively. LAI-Net was trained 10 times, with each training run using a different random seed for sampling from the same generative model.

**Table 2:** Accuracy (Precision in %) of LAI-Net trained on synthetic data that is *X* × larger than real data and evaluated on haptool-admixed-30 test data. The synthetic data was generated by PCA-DDPM (top of table) or Bernoulli model (bottom of table). Ancestry coefficients in test data are 0.16, 0.33, 0.51 for AFR, EAS, EUR respectively. LAI-Net was trained 10 times, with each training run using a different random seed for sampling from the same generative model. Results of the same experiment on haptool-admixed-60 are given in Table S2.

|  | AFR | EAS | EUR | Weighted<br>average |
| --- | --- | --- | --- | --- |
| **Real** | 93.65 ± 0.12 | 94.6 ± 0.14 | 95.67 ± 0.07 | 95.0 ± 0.08 |
| **PCA-DDPM** |  |  |  |  |
| same size (Table 1) | 93.49 ± 0.15 | 93.36 ± 0.16 | 95.02 ± 0.06 | 94.23 ± 0.08 |
| 4x larger | 93.62 ± 0.1 | 93.72 ± 0.14 | 95.12 ± 0.1 | 94.42 ± 0.05 |
| **Bernoulli** |  |  |  |  |
| same size (Table 1) | 84.9 ± 3.21 | 79.33 ± 2.94 | 88.93 ± 1.56 | 85.12 ± 2.27 |
| 2x larger | 84.11 ± 1.7 | 78.78 ± 1.66 | 88.63 ± 0.74 | 84.65 ± 1.16 |
| 4x larger | 83.41 ± 0.71 | 78.03 ± 0.53 | 88.31 ± 0.35 | 84.13 ± 0.44 |
| 6x larger | 84.93 ± 2.71 | 79.81 ± 2.62 | 89.19 ± 1.27 | 85.41 ± 1.93 |

Interestingly, although the baseline random matrix Bernoulli model generates synthetic genomes of notably poor quality (**FIG.S2**), it still holds practical utility by providing a training data substrate that enables relatively good classification performance. This distinction between quality and utility suggests that synthetic genomes can be tailored for specific tasks in applied settings, without placing excessive emphasis on general quality thresholds.

A promising use-case for artificial genomes could be addressing dataset imbalance prevalent in genomic research, particularly due to underrepresented groups. As we demonstrated in this work, artificial genomes can be used as alternatives to real genomes with comparable performance on complex tasks. Here we explored scenarios where the training data for generative models sufficiently covers different modes (i.e., super-populations) but we did not focus on conditional generation on smaller modes such as populations. Modeling the low-density regions of the data space is challenging, as generative models tend to struggle with accurately capturing the distribution of these sparse areas. Future work can explore approaches to overcome this difficulty, potentially by auditing the sampling process with an acceptance criterion to ensure that samples from low-density regions are adequately represented (Alaa et al., 2022).

Overall, this study demonstrates fast and high-quality generation of artificial genomes using a novel compact diffusion model, and confirms their usefulness for local ancestry inference by achieving comparable performance to predictive models trained on real data. This advance opens new avenues for using synthetic data to enhance genomic research, particularly when real data is limited or access is constrained.

## 5 Acknowledgements

This work benefited from Inria TAU computing resources, and funding from ANR-20-CE45-0010-01 RoDAPoG and Human Frontier Science Project (number RGY0075/2019). We thank Jazeps Medina Tretmanis and Fanny Pouyet for insightful discussions.

## 6 Availability of data and materials

All the relevant data and code is publicly available at https://gitlab.inria.fr/ml_genetics/public/artificial_genomes/-/tree/master/1000G_real_genomes for data and https://gitlab.inria.fr/ml_genetics/public/artificial_genomes/-/tree/master/LAI for the models.

## S1 Appendix

### S1.1 Dataset description

**Figure S1:**
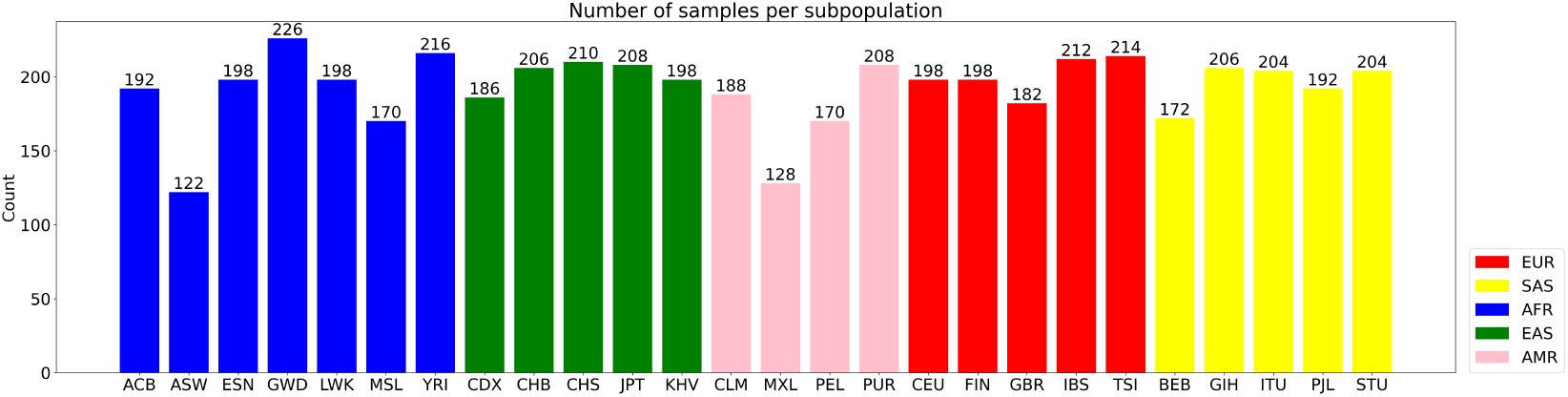
Distribution of the data per population. Population labels refer to intra-continental populations: ACB (African Caribbean in Barbados), ASW (African Ancestry in Southwest US), ESN (Esan in Nigeria), GWD (Gambian in Western Division, The Gambia - Mandinka), LWK (Luhya in Webuye, Kenya), MSL (Mende in Sierra Leone), YRI (Yoruba in Ibadan, Nigeria), CDX (Chinese Dai in Xishuangbanna, China), CHB (Han Chinese in Beijing, China), CHS (Han Chinese South), JPT (Japanese in Tokyo, Japan), KHV (Kinh in Ho Chi Minh City, Vietnam), CLM (Colombian in Medellin, Colombia), MXL (Mexican Ancestry in Los Angeles, California), PEL (Peruvian in Lima, Peru), PUR (Puerto Rican in Puerto Rico), CEU (Utah residents (CEPH) with Northern and Western European ancestry), FIN (Finnish in Finland), GBR (British in England and Scotland), IBR (Iberian populations in Spain), TSI (Toscani in Italy), BEB (Bengali in Bangladesh), GIH (Gujarati Indians in Houston, TX), ITU (Indian Telugu in the UK), PJL (Punjabi in Lahore, Pakistan), STU (Sri Lankan Tamil in the UK).

#### S1.2 Light PCA-DDPM architecture

The denoising neural network has 400K parameters, and achieves great generative abilities at 20K steps, amounting to a training time (wall clock) of roughly 10 minutes on a single A100-40GB GPU. The number of diffusion steps is set to 1,000. Sampling the DDPM to generate thousands of samples takes only few seconds.

- **Label embedding layer**: `nn.Embedding(3, 3)`
- **Time encoding layer**: `SinusoidalTimeEmbedding`
- **Predictor**:
  - **Input layer**: concatenation of 6 features and 3 label features
  - **Linear input layer**: 256 neurons
  - **1st residual block**:
    * Linear transformation with 256 neurons
    * Time integration via linear transformation with 256 neurons
    * Linear transformation with 256 neurons
    * Residual connection added
  - **2nd residual block**:
    * Linear transformation with 256 neurons
    * Time integration via linear transformation with 256 neurons
    * Linear transformation with 256 neurons
    * Residual connection added
  - **Output layer**: 6 neurons
- **Forward pass**:
  - Input tensor *x*_*t*_, time tensor, and super-population label processed through:
    * Embedding layer to obtain label representation
    * Concatenation of *x*_*t*_ and label embedding
    * Linear transformation to the input layer
    * Activation function applied to each layer
    * Residual connections added after each block
    * Final output layer produces 6 outputs
- **Optimizer**: `Adam` with initial learning rate 0.0003
- **Learning rate scheduler**: `CosineAnnealingWarmRestarts`
- **Warmup scheduler**: `LinearWarmup` with 1000 steps warmup period

**Table S1:** Training GPU & runtime comparison between models for 65K SNP dataset.

| Models | Params | GPUs | Wall clock |
| --- | --- | --- | --- |
| Light PCA-WGAN with NN | 750K | 1-A100-40GB | ~2 hours |
| Light PCA-DDPM | 400K | 1-A100-40GB | ~10 minutes |

#### S1.3 Relation between number of generations and window size

*Haptools* (Massarat et al., 2023) simulates admixed samples given ancestral source genomes, ancestry coefficients, number of generations over which the forward simulation will run and a genetic map. Below we explain the relationship between generations and expected size of the ancestral genomic segments.

Let *n* be the number of generations, *ρ* the recombination rate, *ω* the size of the window (in base-pairs, abbreviated bp) defined by crossover events. For each position in a genome, at each generation, there is a probability *ρ* of recombination occurring at that site. The probability of having at least one recombination at a single position during *n* generation is 1 − (1 − *ρ*)^*n*^, which as ρ ∽ 10^−8^ can be approximated by *nρ*. We can now model recombination as a Bernoulli trial, where the distance *ω* between successive recombination events follows a geometric law of parameter *p* = 1 − (1 − *ρ*)^*n*^∽ *nρ*. The expected distance between recombination events is given by

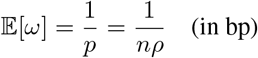

Hence

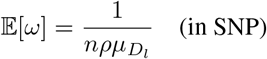

where 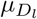 is the average distance between SNP in chromosome *l*, in bp unit and *ρ* can be estimated from a genetic map. For the 65,535 SNP data spanning chr1:534247-81813279, we estimated *ρ* = 2.3 *×* 10^−8^ and 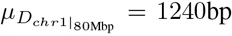. For *n* = 30, we have 𝔼[*ω*] = 6183 SNPs, *i*.*e*., the average window size defined by a crossover events is *>*∽12 larger than LAI-Net resolution set to 500 SNPs. Note that a crossover event can link together segments of the same ancestry, however we are only interested here to crossover events corresponding to changes in ancestry meaning that actual average window size will be higher. Precisely, let *a* be an ancestry of proportion *x*_*a*_ in the population. At a recombination event, any segment then has probability (1 − *x*_*a*_) to recombine with a segment of an ancestry different of *a*. Therefore the average length of a segment of ancestry *a* will be

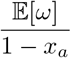

In our case it means, for 30 generations, an average segment length of 7361 SNPs for AFR, 9228 SNPs for EAS and 12618 SNPs for EUR.

#### S1.4 High quality artificial genomes

**Figure S2:**
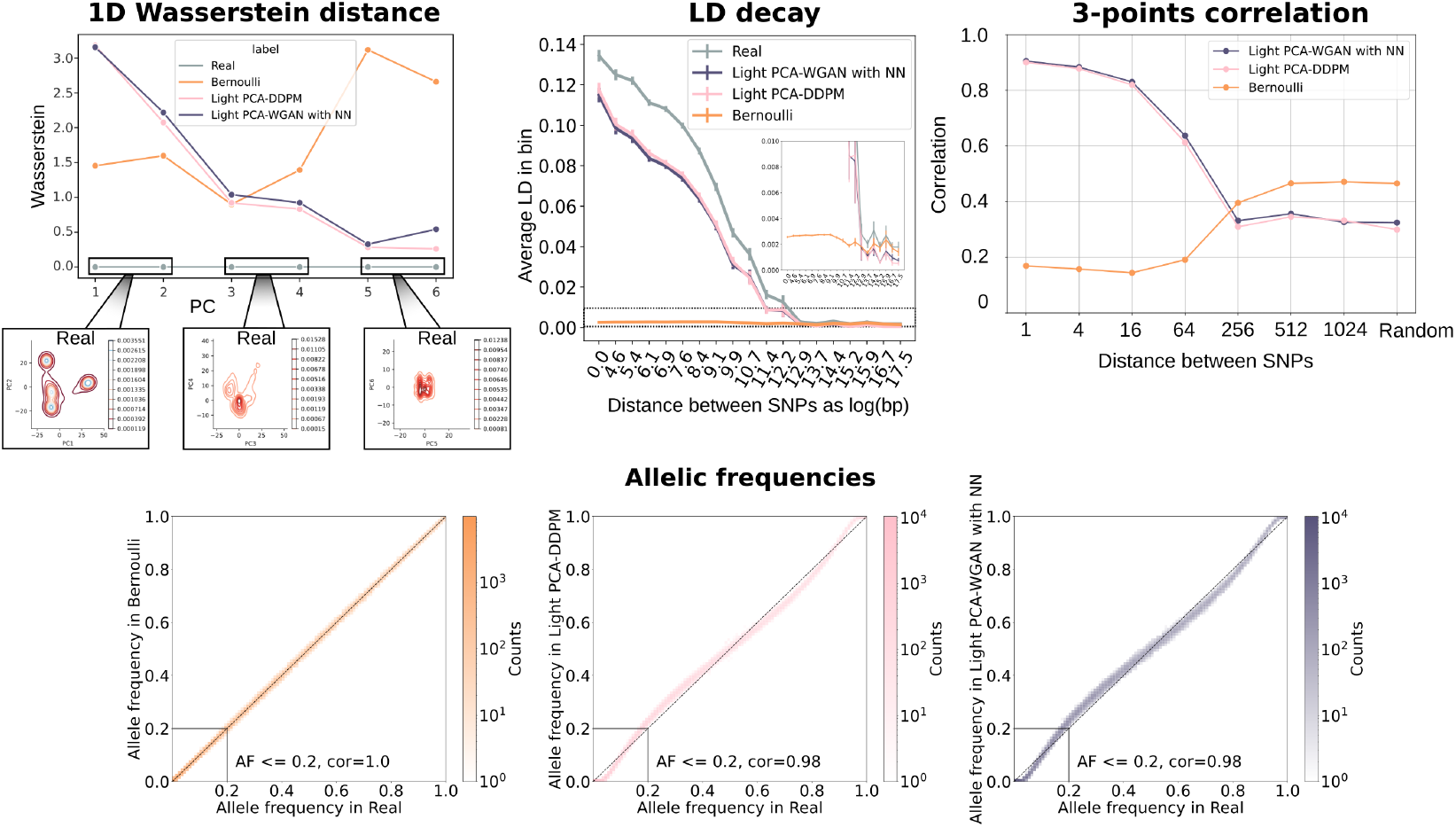
Population genetics summary statistics. The real and the synthetic data were concatenated into a single dataset, to which PCA is applied. **Upper left**. The 1D Wasserstein distance is computed on each PCs. Real data is the grey curve, Bernoulli in orange, Light PCA-DDPM in pink and Light PCA-WGAN from (Yelmen et al., 2023) in dark purple. The density plot of real data for the first six PCs is shown. **Upper middle**. LD decay approximation. The inset is a zoom on small LD values. **Upper right**. Three-points correlation (3pt-corr) for triplets separated by a varying amount of SNPs, displayed as Corr( 3pt-corr[synthetic data], 3pt-corr[real data] ). **Lower middle**. 2D Histogram of the allelic frequencies in real data (x-axis) and in AGs produced by the models.

**Figure S3:**
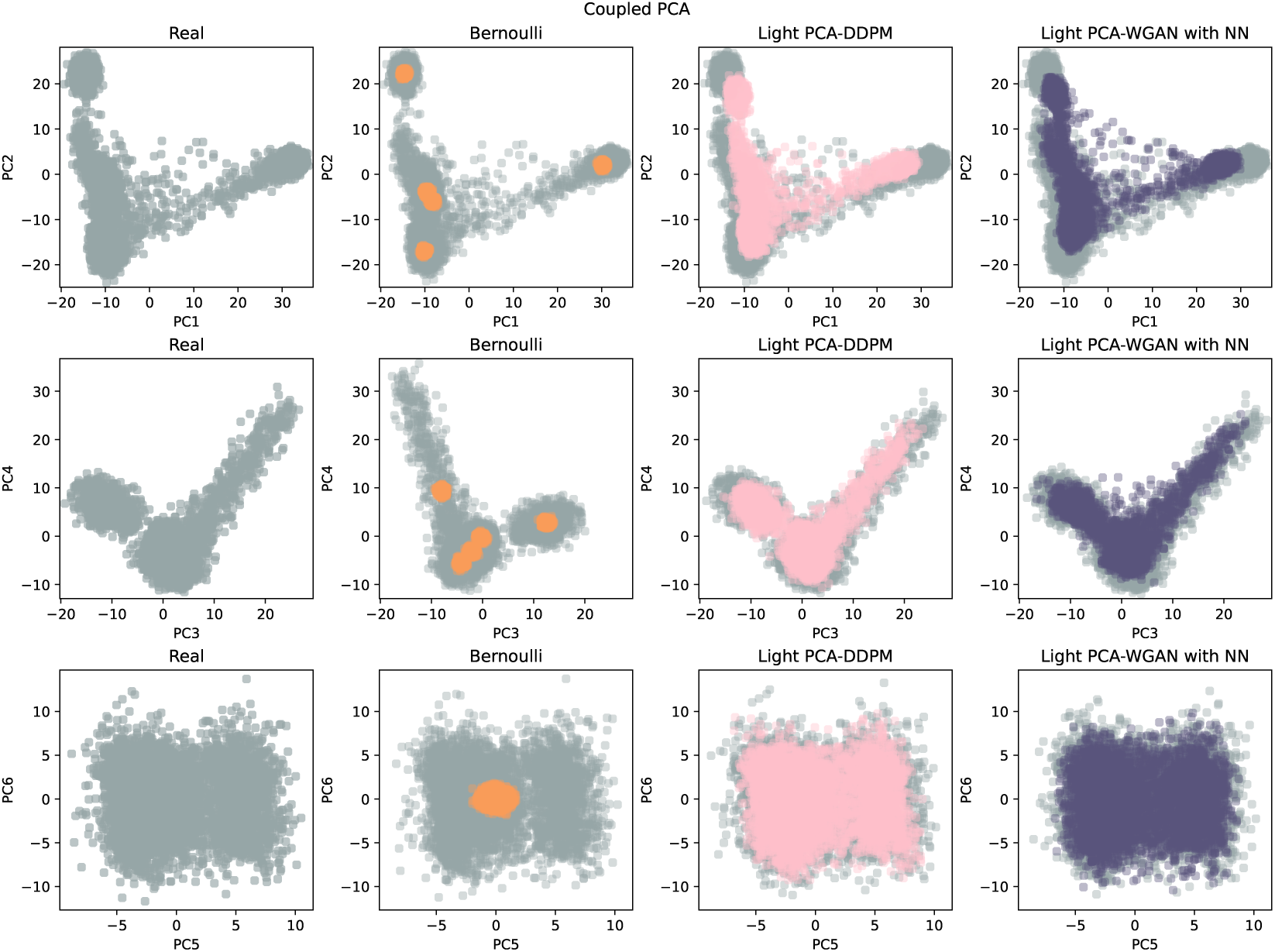
Coupled PCA. Real and synthetic samples are concatenated into a single dataset upon which PCA is applied. Synthetic samples are generated by Bernoulli (orange), Light PCA-DDPM (pink), Light PCA-WGAN (dark purple). First row corresponds to (PC1,PC2), second to (PC3,PC4) and third to (PC5,PC6).

#### S1.5 Increasing synthetic sample size

**Table S2:** Accuracy (Precision in %) of LAI-Net trained on synthetic data that is *X* larger than real data and evaluated on haptool-admixed-60 test dataset. The synthetic data was generated by PCA-DDPM (top of table) or Bernoulli model (bottom of table) . Ancestry coefficients in test data are 0.16, 0.33, 0.51 for AFR, EAS, EUR respectively. LAI-Net was trained 10 times, with each training run using a different random seed for sampling from the same generative model.

|  | AFR | EAS | EUR | Weighted<br>average |
| --- | --- | --- | --- | --- |
| **Real** | $88.29 \pm 0.19$ | $90.99 \pm 0.12$ | $92.94 \pm 0.08$ | $91.55 \pm 0.06$ |
| **PCA-DDPM** |  |  |  |  |
| same size (Table 1) | $88.57 \pm 0.34$ | $89.78 \pm 0.15$ | $92.25 \pm 0.11$ | $90.85 \pm 0.09$ |
| 4x larger | $88.56 \pm 0.21$ | $90.31 \pm 0.18$ | $92.38 \pm 0.12$ | $91.08 \pm 0.04$ |
| **Bernoulli** |  |  |  |  |
| same size (Table 1) | $79.0 \pm 3.4$ | $76.25 \pm 2.56$ | $87.19 \pm 1.02$ | $82.27 \pm 1.9$ |
| 2x larger | $78.17 \pm 1.81$ | $75.7 \pm 1.41$ | $87.02 \pm 0.48$ | $81.87 \pm 0.96$ |
| 4x larger | $77.55 \pm 0.84$ | $75.25 \pm 0.39$ | $86.93 \pm 0.23$ | $81.57 \pm 0.34$ |
| 6x larger | $78.98 \pm 2.9$ | $76.66 \pm 2.32$ | $87.46 \pm 0.84$ | $82.54 \pm 1.64$ |

